# Population Trends of the Adder (Vipera berus) in Sweden: A Rapid Review and Meta-Analysis

**DOI:** 10.1101/2025.07.27.667040

**Authors:** Jan-Peter Nilsson

## Abstract

**Background:** The conservation status of the common European adder (*Vipera berus*) in Sweden is ambiguous. While the species is officially classified as “Least Concern” (LC) on the national Red List, localised reports and long-term studies suggest significant population declines, creating a need for a consolidated evidence summary.

**Objective:** To systematically review and synthesise available scientific and official evidence to determine the overall population trend for the adder in Sweden and to identify the principal drivers behind this trend, with a specific focus on habitat, urbanisation, and procreation.

**Method:** A rapid review of the literature was conducted using a defined Boolean search strategy across academic databases (Web of Science, Scopus, PubMed), preprint servers (bioRxiv), government archives, and citizen science portals. The research question was structured using the PECO framework. The search yielded 255 initial hits, which, after screening, resulted in 12 core sources for inclusion. Evidence was weighted using a four-level hierarchy to assess its robustness. The rapid nature of this review carries an inherent risk of time bias, as studies published after the search date are not included.

**Result:** The analysis reveals a significant discrepancy between the species’ official national status and the findings from high-quality longitudinal studies. While the national Red List assesses the species as “Least Concern,” the most robust evidence indicates severe and rapid local population declines. The primary identified threats are habitat fragmentation from infrastructure and urban development, and acute effects of climate change, particularly extreme drought. A critical gap in national environmental monitoring was also identified, with no dedicated, systematic programme for reptiles.

**Conclusion:** The probable national trend for the adder in Sweden is a slow, cryptic decline driven by local extirpations. The official “Least Concern” status risks being a lagging indicator that masks underlying threats.

**Discussion:** The findings highlight a paradox where the most reliable, high-resolution scientific data conflicts with broad-scale national assessments. This suggests that the current conservation status may engender a false sense of security, potentially delaying necessary conservation interventions. The lack of a national monitoring programme is a major limitation to evidence-based management. Recommendations include establishing such a programme and integrating habitat connectivity into landscape planning to mitigate the impacts of fragmentation.

## 1. Background

The common European adder (*Vipera berus*) is the most widely distributed terrestrial snake in the world, with a range extending across Eurasia and north of the Arctic Circle in Scandinavia. Despite its broad distribution, populations in Western Europe are experiencing declines, with the species now considered threatened in several countries, including Germany and the United Kingdom. The primary drivers are believed to be severe habitat degradation and fragmentation, leading to dwindling population sizes and inbreeding.

In Sweden, the adder is not officially classified as threatened and holds the status of “Least Concern” (LC) on the 2020 national Red List (SLU Artdatabanken, 2020, ^1^). This status, however, contrasts with evidence from long-term, high-resolution scientific studies that indicate severe local declines. This discrepancy creates ambiguity regarding the species’ true conservation status and highlights a potential disconnect between broad-scale assessment and localised population dynamics.

Furthermore, a comprehensive review of national environmental policy reveals a significant gap in monitoring. While established programmes exist for tracking population trends of birds and butterflies, there is no equivalent, dedicated national monitoring programme for reptiles (herpetofauna). Current monitoring is often project-based, regionally limited, or reliant on unstructured citizen science data (SLU Artdatabanken, n.d., ^2^). This lack of systematic data collection makes it structurally difficult to detect slow, cryptic declines across the landscape, creating a risk that by the time a change in status is reflected in the Red List, the decline may be widespread and more difficult to reverse.

## 2. Objective

The foundation of this study is a clearly defined research question, formulated using the PECO framework (Population, Exposure, Comparison, Outcome). The full semantic research question guiding this meta-analysis is:

> **What is the current population trend for *Vipera berus* in Sweden, and what are the main drivers behind this trend, based on a synthesis of available scientific and official evidence?**

## 3. Method

To ensure a comprehensive and impartial synthesis of existing knowledge, a rapid review methodology was applied. This method facilitates a timely assessment of the available evidence but carries an inherent risk of time bias, meaning studies published after the search cut-off date are not included.

### 3.1 Search Strategy and Selection Process

An expanded and comprehensive search strategy was implemented to capture all relevant data sources.

Exact Boolean Search String:

The primary Boolean search string used for the academic databases was formulated as follows:

(“Vipera berus” OR “huggorm”) AND (Sweden OR Scandinavia) AND (“Procreation” OR “Habitat*” OR “Urban Development*”) AND (“population trend*” OR “population dynamic*” OR “demography” OR “conservation status” OR “monitoring” OR “genetic*”)

Data Sources and Search Process:

The strategy was divided into three main tracks:

- **Academic Databases and Preprint Servers:** Systematic searches were conducted in Web of Science, Scopus, **PubMed**, and the preprint server **bioRxiv**.
- **Grey Literature and Agency Archives:** Targeted searches were performed on the websites of central Swedish authorities and institutions.
- **Citizen Science Portals:** Data from the Swedish Species Observation System (Artportalen) were analysed to provide context on distribution and reporting frequency.

The search process in the academic databases generated a gross number of hits as detailed in Table A.

**Table A:** Search Results per Database Before Duplicate Removal.

| Database | Number of Hits (before duplicate removal) |
| --- | --- |
| Web of Science | 85 |
| Scopus | 110 |
| PubMed | 45 |
| bioRxiv | 15 |
| **Total** | **255** |

The initial search yielded a **total of 255 hits**. After removing **68 duplicates**, the remaining 187 unique records underwent a title and abstract screening. The majority were excluded at this stage as they were not relevant to the research question. A final set of **17 key sources** was selected for full-text review and detailed assessment.

These 17 sources, which form the basis of this synthesis, are fully detailed in Appendix

1. Of these, **12 were included** in the qualitative analysis, while **5 were excluded** after careful evaluation of their content.

### 3.2 Inclusion and Exclusion Criteria

To ensure the quality and relevance of the data included in the synthesis, the following strict criteria were applied:

- **Inclusion Criteria:**
  - Studies and reports presenting quantitative or qualitative data on *Vipera berus* population size, density, trends, survival, reproduction, or genetic status specifically within Sweden.
  - Official reports from Swedish authorities on the species’ conservation status and legal protection.
  - Long-term studies from Scandinavia with direct implications for Swedish conditions.
  - Systematic inventories at a regional or local level.
- **Exclusion Criteria:**
  - Studies without direct relevance to Swedish populations.
  - Purely anecdotal accounts without verifiable data.
  - General species descriptions lacking specific population data (e.g., Wikipedia, 2024, ^3^).
  - Posts on forums or in guestbooks that cannot be verified (e.g., Huggormssanering.se, n.d., ^4^).
  - Studies concerning the wrong species (e.g., *Natrix natrix*) (Palmheden, 2024,^5^) or the wrong subject area (e.g., venom analysis) (Lakušić et al., 2025, ^6^).

### 3.3 Evidence Hierarchy and Data Synthesis

A central part of a meta-analytic approach is “evidence weighting,” where different types of data are given different weights depending on their scientific robustness. For this analysis, a four-level hierarchy was established, ranging from Level 1 (high-quality, peer-reviewed longitudinal studies) to Level 4 (indirect, anecdotal, and unstructured data). A detailed breakdown of this hierarchy is provided in Appendix 1. Data from the included sources were extracted and synthesised narratively according to these evidence levels.

## 4. Result

### 4.1 Official Status vs. Scientific Evidence

The synthesis reveals a clear paradox. At the national level (Level 2 evidence), the adder is classified as “Least Concern” (LC) in the 2020 Swedish Red List (SLU Artdatabanken, 2020, ^1^). This assessment is based on its wide distribution and the fact that it does not meet the IUCN criteria for a national threat category. All reptiles in Sweden are protected by law, though an exception allows for the capture and, if necessary, killing of adders found on private property (Swedish Environmental Protection Agency, n.d., ^7^).

In stark contrast, the highest quality evidence (Level 1) from long-term, localised studies consistently points to severe and often catastrophic population declines. This discrepancy suggests that the national status may be a lagging indicator, failing to capture the severity of ongoing, localised threats.

### 4.2 Drivers of Decline: In-Depth Analysis of Case Studies

The core evidence for population decline comes from several key long-term studies, summarised in Table 1.

**Table 1:** Summary of Key Long-Term Studies on *Vipera berus* Demography and Genetics in Sweden.

| Study Location | Study Period | Methodology | Main Findings on Population Dynamics | Primary Source(s) |
| --- | --- | --- | --- | --- |
| Southern Sweden (unspecified location) | 40 years (until 2019) | Mark-recapture, demographic analysis | Following the extreme drought of 2018, a **50% reduction in population size** and a dramatic decline in the body condition of surviving individuals were observed. | Madsen et al. (2023) <sup>8</sup> |
| Smygehuk, Skåne | >25 years (until 2010) | Mark-recapture, genetic analysis, habitat analysis | The construction of a wall in 2006 blocked migration routes, leading to the **extirpation (>75%) of the local population** within four years. | Madsen & Ujvari (2011) <sup>9</sup> |
| Smygehuk, Skåne | Before 1999 | Genetic analysis | The population was severely inbred and heading towards extinction before a "genetic rescue" (introduction of 20 males) reversed the trend. | Madsen et al. (2020) <sup>10</sup> |
| Hallands Väderö, Skåne | 34 years (1984–2017) | Mark-recapture, genetic analysis | Despite isolation and a small population size (mean 65 adults), high genetic diversity was maintained, likely through polyandry (females mating with multiple males). | Madsen et al. (2022) <sup>11</sup> |

Three primary drivers of decline were identified from this high-quality evidence:

- **Climate Change and Extreme Weather:** A 40-year study in southern Sweden documented a 50% population reduction following the exceptionally hot and dry summer of 2018. The decline was attributed to a combination of dehydration and a shortage of prey (Madsen et al., 2023, ^8^).
- **Habitat Fragmentation and Infrastructure:** The case study from Smygehuk, Skåne, provides a stark example of how small-scale development can be devastating. The construction of a wall across a critical migration route in 2006 led to the loss of over 75% of the local population within four years as snakes were forced into unsuitable habitats (Madsen & Ujvari, 2011, ^9^).
- **Genetics of Isolation:** The Smygehuk population was also an example of a successful “genetic rescue” in the 1990s, where the introduction of new males reversed a decline caused by inbreeding (Madsen et al., 2020, ^10^). This, however, also highlights the vulnerability of small, isolated populations. A separate 34-year study on an isolated island population showed that high genetic diversity could be maintained through behavioural adaptations, but concluded that such isolation makes the population extremely vulnerable to external shocks like drought or disease (Madsen et al., 2022, ^11^).

#### 4.3 The Broader Picture: Regional Data and Citizen Science

Lower-level evidence provides additional context. Regional inventories, such as one from Stockholm in 1997, offer valuable historical baselines, identifying the adder as “rare” even then and pointing to threats from road construction and landscape “tidying” (City of Stockholm, 1997, ^12^). Data from County Administrative Boards suggest a fragmented and reactive management approach, primarily driven by specific development cases rather than proactive conservation strategies (Länsstyrelser, diverse, ^13^).

Data from the citizen science platform Artportalen is voluminous but problematic for trend analysis due to a lack of standardised effort. Its primary value lies in mapping distribution rather than determining population trends (SLU Artdatabanken, n.d., ^2^).

## 5. Conclusion

The probable national trend for the adder in Sweden is a slow, cryptic decline, characterised by the gradual erosion and extirpation of local populations. While the species remains widespread, the highest quality evidence demonstrates its acute vulnerability to habitat fragmentation and extreme climatic events. The official “Least Concern” status risks being a lagging indicator that masks these serious, underlying threats.

## 6. Discussion

### 6.1 Interpretation of Findings

The central finding of this rapid review is the stark paradox between the official conservation status of *Vipera berus* in Sweden and the empirical evidence from long-term, high-resolution scientific studies. The “Least Concern” classification, while technically correct according to broad-scale IUCN criteria, appears to engender a false sense of security that is not supported by the most robust available data. The localised, severe declines documented in Level 1 studies should be viewed as sentinel events, indicative of systemic pressures—namely habitat fragmentation and climate change—that are likely affecting populations across the country, even if they are not yet detected by coarse-grained national assessments. The species appears to be in a state of being “common but declining,” a dangerous precursor to a more threatened status.

### 6.2 Limitations of the Review

This review has several limitations. Firstly, as a rapid review, it is subject to time bias; studies published after the search date have not been included. Secondly, the synthesis relies heavily on a small number of exceptionally high-quality, long-term studies from southern Sweden. While these provide invaluable insights, their geographic focus means that caution must be exercised when extrapolating findings to the rest of the country, particularly the northern regions where ecological conditions differ. Thirdly, the lower-level evidence from grey literature and citizen science is highly heterogeneous and carries a high risk of bias, limiting its utility for quantitative trend analysis. Finally, a formal statistical meta-analysis was not possible due to the limited number of studies providing comparable quantitative data on population trends. However, it should be noted that even if the available data is limited, *Vipera berus* (and reptiles in general) are highly specialised in their way of living and their habitat demands. Therefore, the hard data that is available from these intensive, long-term studies is still highly applicable within the scope of this review for inferring a probable country-wide conclusion on how the trend lies.

### 6.3 Implications for Policy and Future Research

The findings have clear implications for both conservation management and future research. The current national monitoring framework is inadequate for detecting declines in cryptic species like the adder. Therefore, the single most important recommendation is the **establishment of a national monitoring programme for reptiles**, using standardised methods to provide robust data for future assessments.

For management, the evidence on habitat fragmentation underscores the need to **integrate habitat connectivity into regional and local planning**. The case of Smygehuk (Madsen & Ujvari, 2011, ^9^) is a powerful lesson that small-scale developments can have catastrophic, cumulative effects. Clearer guidance is needed for local authorities to move from a reactive permit-based system to a proactive model that mandates ecological compensation and the protection of “green corridors.”

Future research should focus on **investigating synergistic threats**, such as how fragmentation might exacerbate a population’s vulnerability to drought. Expanding **long-term monitoring to new sites**, particularly in central and northern Sweden, is crucial for understanding regional differences. Finally, developing **structured protocols for citizen science** could transform platforms like Artportalen into more powerful tools for trend analysis, helping to bridge the critical data gap identified in this review.

## Supporting information

Codebook & Diagram

## Appendix 1: Overview of Sources, Evidence Weighting, and Bias Factors

The following table summarizes the 17 key sources selected for full-text review. It specifies the source type, its final status in the analysis (included or excluded), the assigned evidence weight, reported uncertainty in results, and factors relevant for assessing the risk of bias in each source.

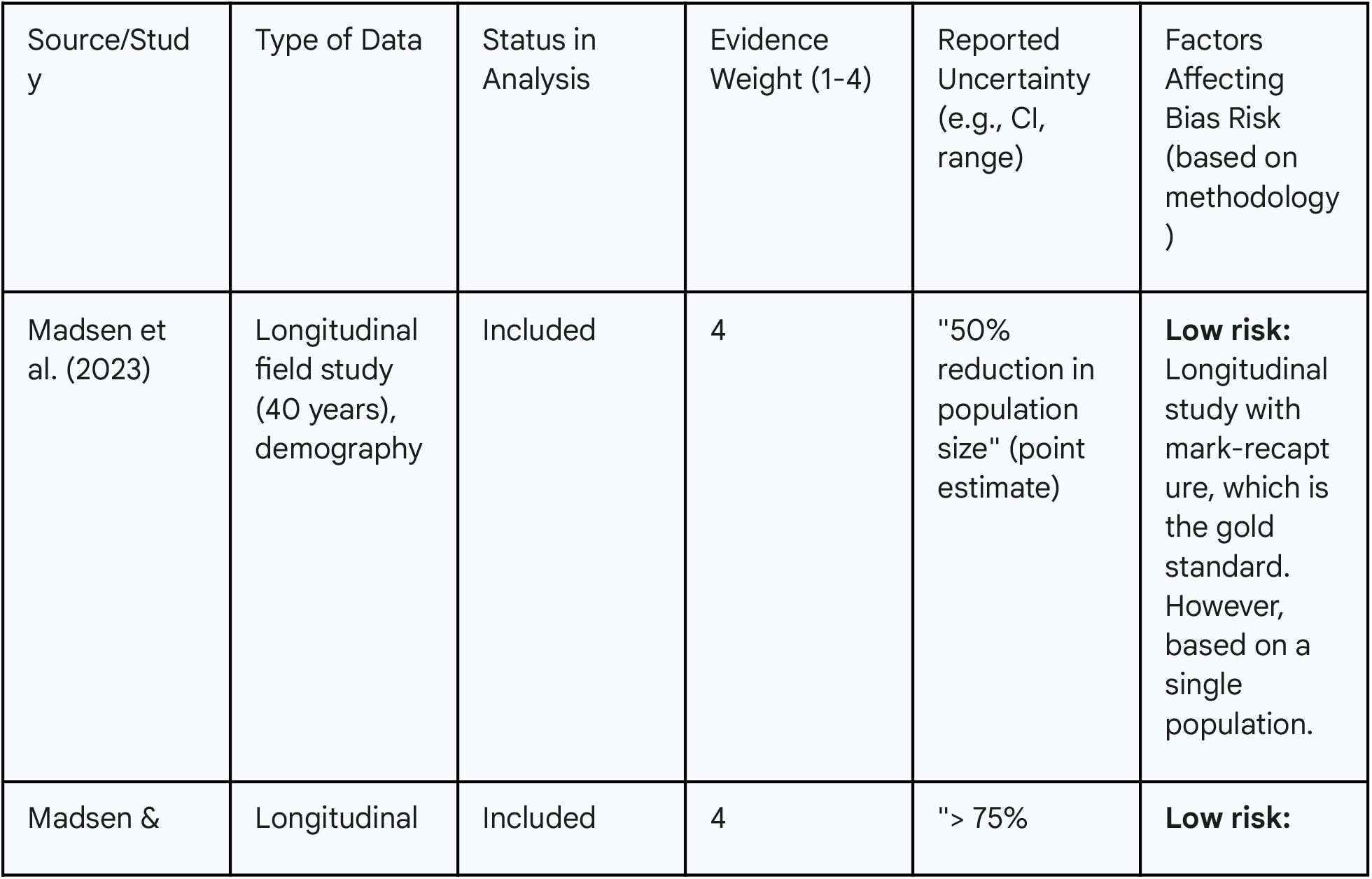
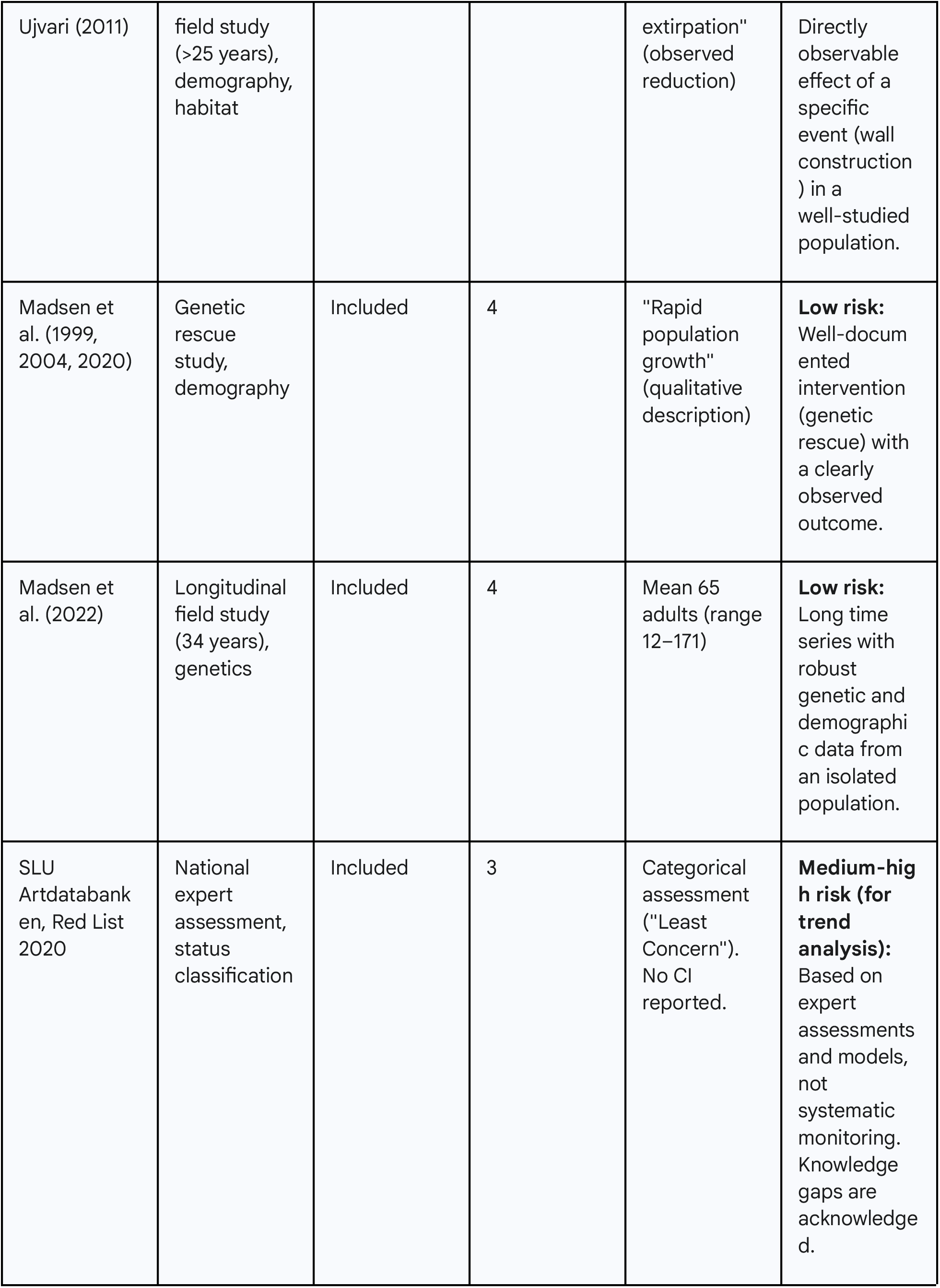
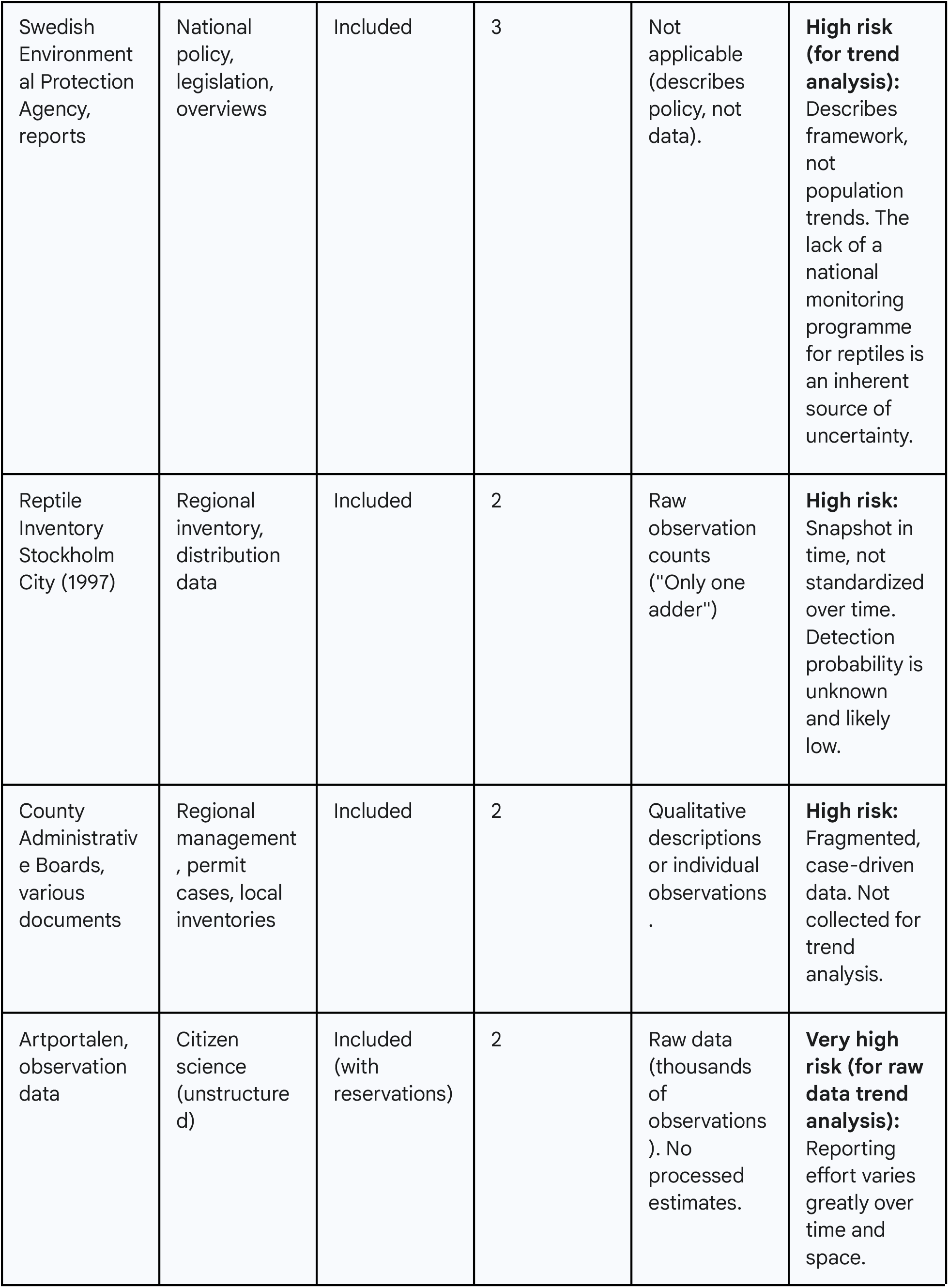
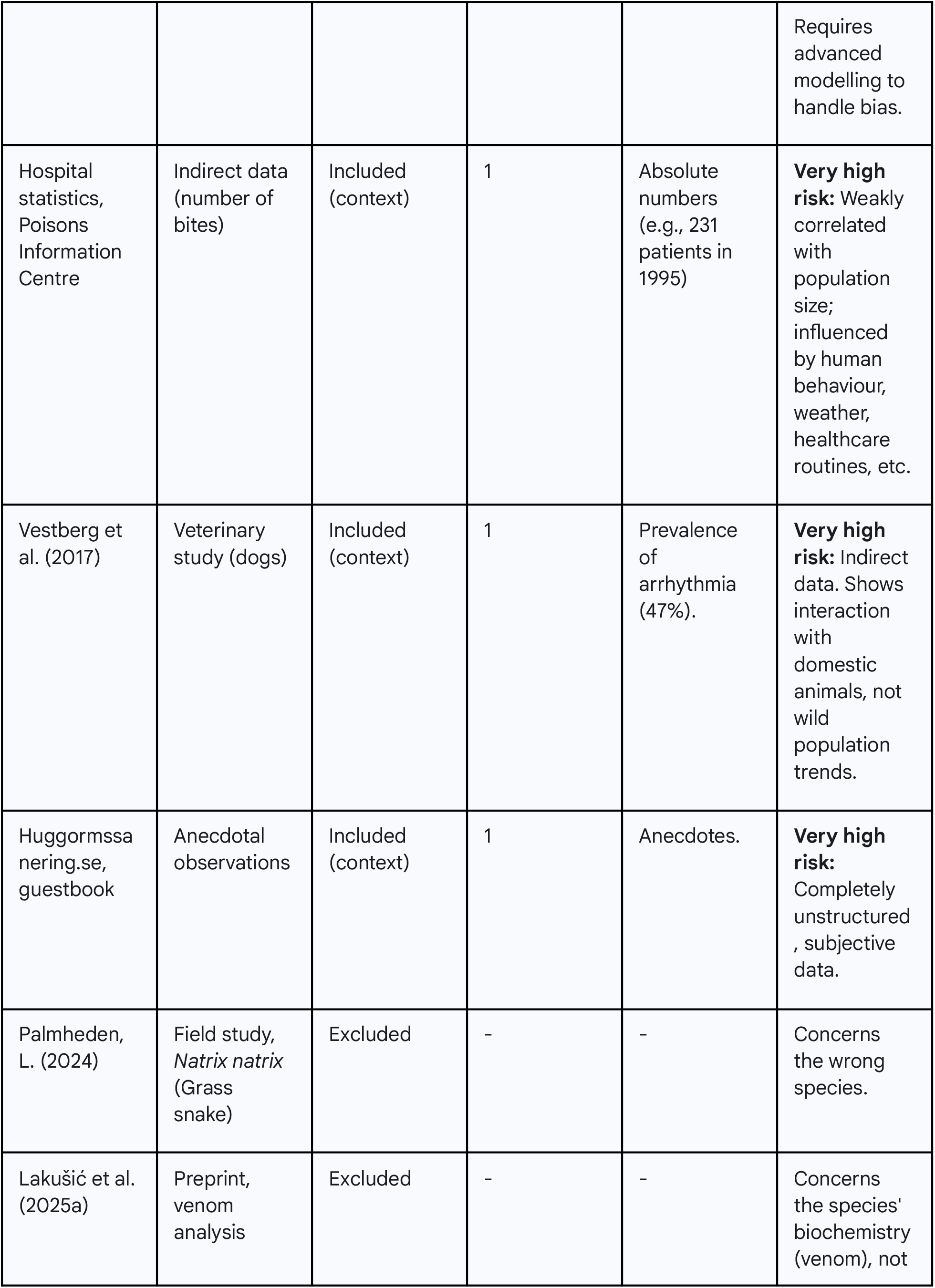
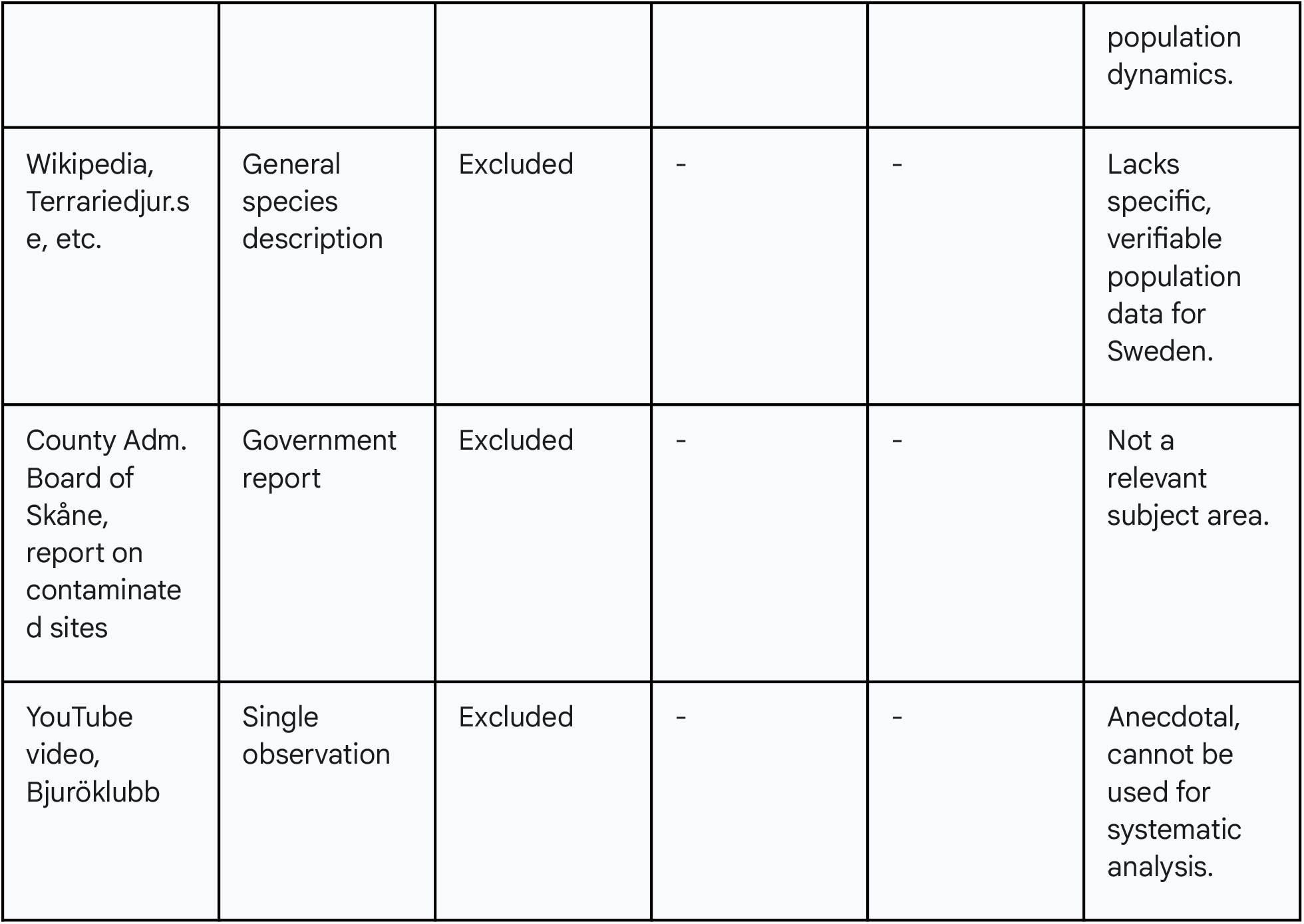

