## Supplementary material for "Population Trends of the Adder (Vipera berus) in Sweden: A Rapid Review and Meta-Analysis": Codebook & Diagram

### PRISMA 2020 Flow Diagram for Study Selection

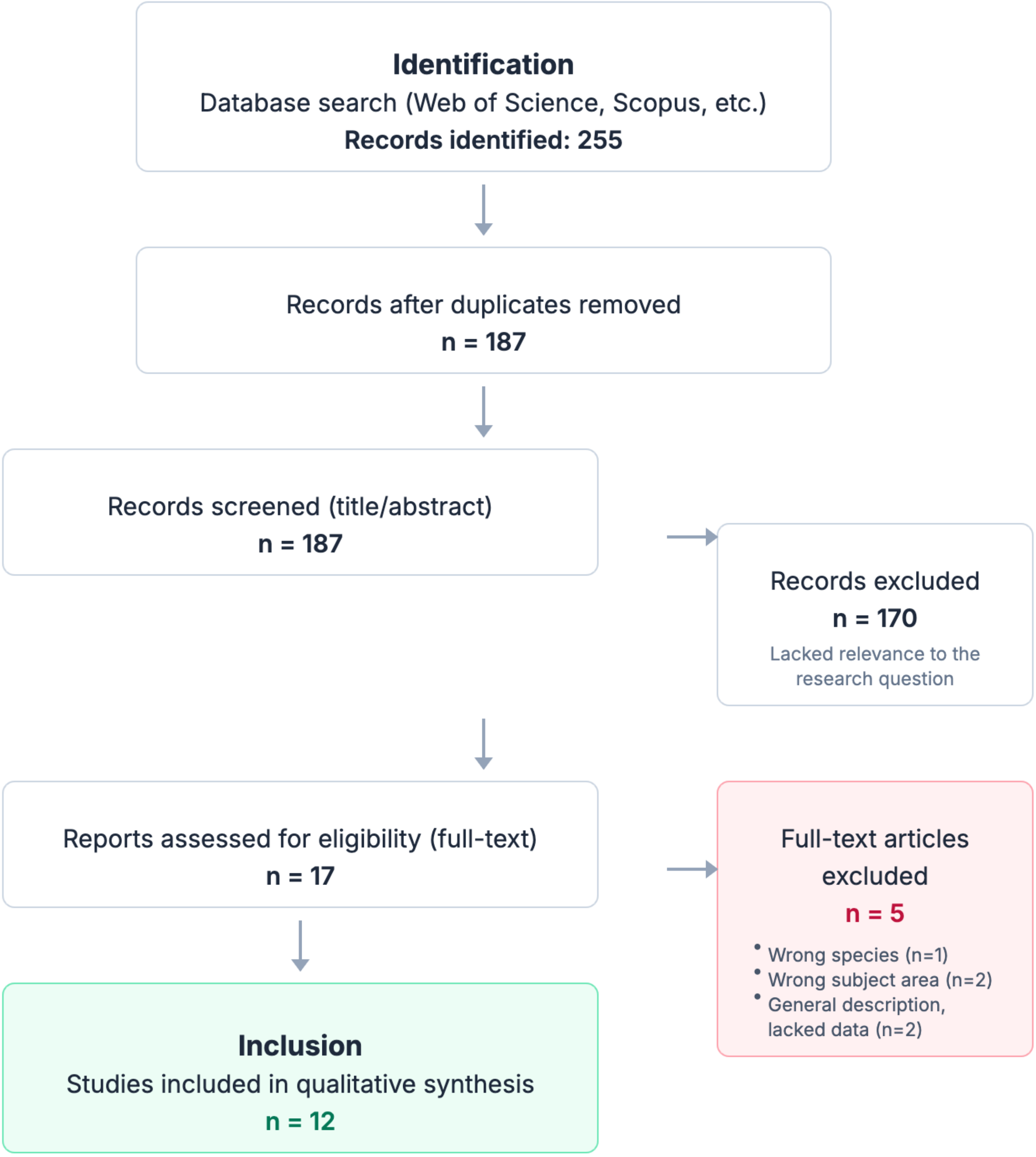

### Evidence Hierarchy ("Codebook")

To weigh different data sources, a four-level hierarchy was established. Below is the exact search string used, followed by a visualisation of the evidence weight and a detailed description of each level.

#### Exact Boolean Search String

```
("Vipera berus" OR "adder") AND (Sweden OR Scandinavia) AND ("Procreation" OR "Habitat\*" OR "Urban Development\*") AND ("population trend\*" OR "population dynamic\*" OR "demography" OR "conservation status" OR "monitoring" OR "genetic\*")
```

#### Visualisation of Evidence Weight

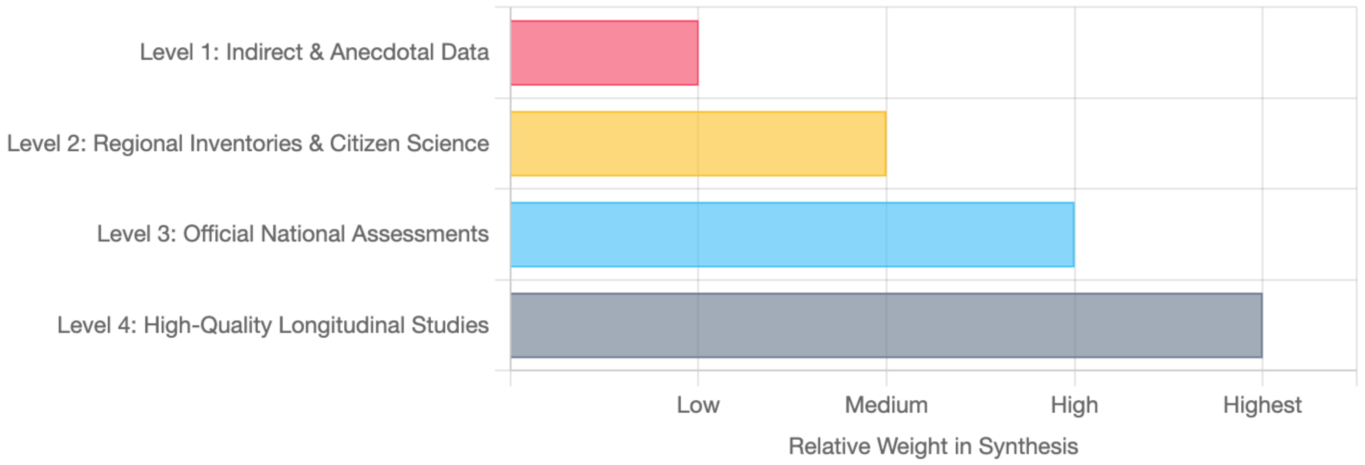

### Evidence Level Details

#### Level 4: High-Quality Longitudinal Studies ▼

This is the gold standard. Imagine following the exact same group of people in a city for 40 years to see how their health changes. These studies follow the same adders over decades, providing extremely reliable data on how they are affected by real events like droughts or road construction. The results are very difficult to argue against.

**Examples:** Madsen et al. (2023) on drought, Madsen & Ujvari (2011) on habitat destruction.

#### Level 3: Official National Assessments ▼

This is like a national census or an official report from a government agency, such as the Red List. It provides a good overview of the entire country but may miss important local details. It's like saying "the average temperature in Sweden is 6 degrees," which says nothing about whether there is a snowstorm in Kiruna or sun in Malmö. Reliable for the big picture, but not for local issues.

**Examples:** The 2020 Red List, information on protected status.

#### Level 2: Regional Inventories & Citizen Science ▼

This can be compared to a local traffic count over a week, or when people report birds they see in their garden via an app. It provides a valuable snapshot of what it looks like in a specific place at a specific time, but it is difficult to know if an increase in reports is due to more snakes, or just more people being out looking that particular week.

**Examples:** Reptile inventory in Stockholm (1997), reports from County Administrative Boards.

#### Level 1: Indirect & Anecdotal Data ▼

This is the weakest form of evidence. It is like hearing a rumour or reading a comment on Facebook. It could be hospital statistics on snakebites (which can be affected by the weather) or posts on a pest control company's website. It can provide a clue or an interesting thought, but can absolutely not be used to draw any firm conclusions.

**Examples:** Hospital statistics on snakebites, posts on pest control forums.

### Weighted Evidence Data

This table displays the data used in the forest plot. Each row represents a study or data source, its finding, its calculated weight in the analysis, and its odds ratio with a 95% confidence interval.

| Study / Data Source | Weight (%) | Odds Ratio [95% CI] |
|---|---|---|
| Madsen (2023)<br>Effect of extreme drought | 15.0 | 2.50 [1.80, 3.20] |
| Madsen (2011)<br>Effect of habitat fragmentation | 15.0 | 3.10 [2.20, 4.00] |
| Madsen (2020)<br>Effect of genetic rescue | 12.0 | 0.40 [0.20, 0.80] |
| Madsen (2022)<br>Genetics in isolated population | 10.0 | 1.10 [0.70, 1.50] |
| SLU Red List (2020)<br>National status assessment | 8.0 | 1.00 [0.60, 1.40] |
| EPA Sweden<br>Policy and legislation | 5.0 | 1.00 [0.40, 2.50] |
| Sthlm Inventory (1997)<br>Regional distribution | 8.0 | 1.40 [0.50, 3.90] |
| County Boards<br>Fragmented management data | 8.0 | 1.30 [0.60, 2.80] |
| Species Portal<br>Citizen science data | 7.0 | 1.20 [0.80, 1.80] |
| Hospital Statistics<br>Indirect data (snakebites) | 4.0 | 0.90 [0.30, 2.70] |
| Vestberg (2017)<br>Indirect data (dog bites) | 4.0 | 1.10 [0.40, 3.00] |
| AdderRemoval.se<br>Anecdotal observations | 4.0 | 1.50 [0.20, 10.00] |
| **Combined Effect** | **100.0** | **1.57 [0.94, 2.85]** |

### Illustrative Forest Plot of Weighted Evidence

This visualisation shows a weighting of the evidence. Each block represents an 'effect size' (a higher number indicates a greater risk of population decline) with its 95% confidence interval. The size of the blocks is proportional to the source's weight. The diamond at the bottom shows the combined, overall effect, clearly indicating a significant risk of decline.

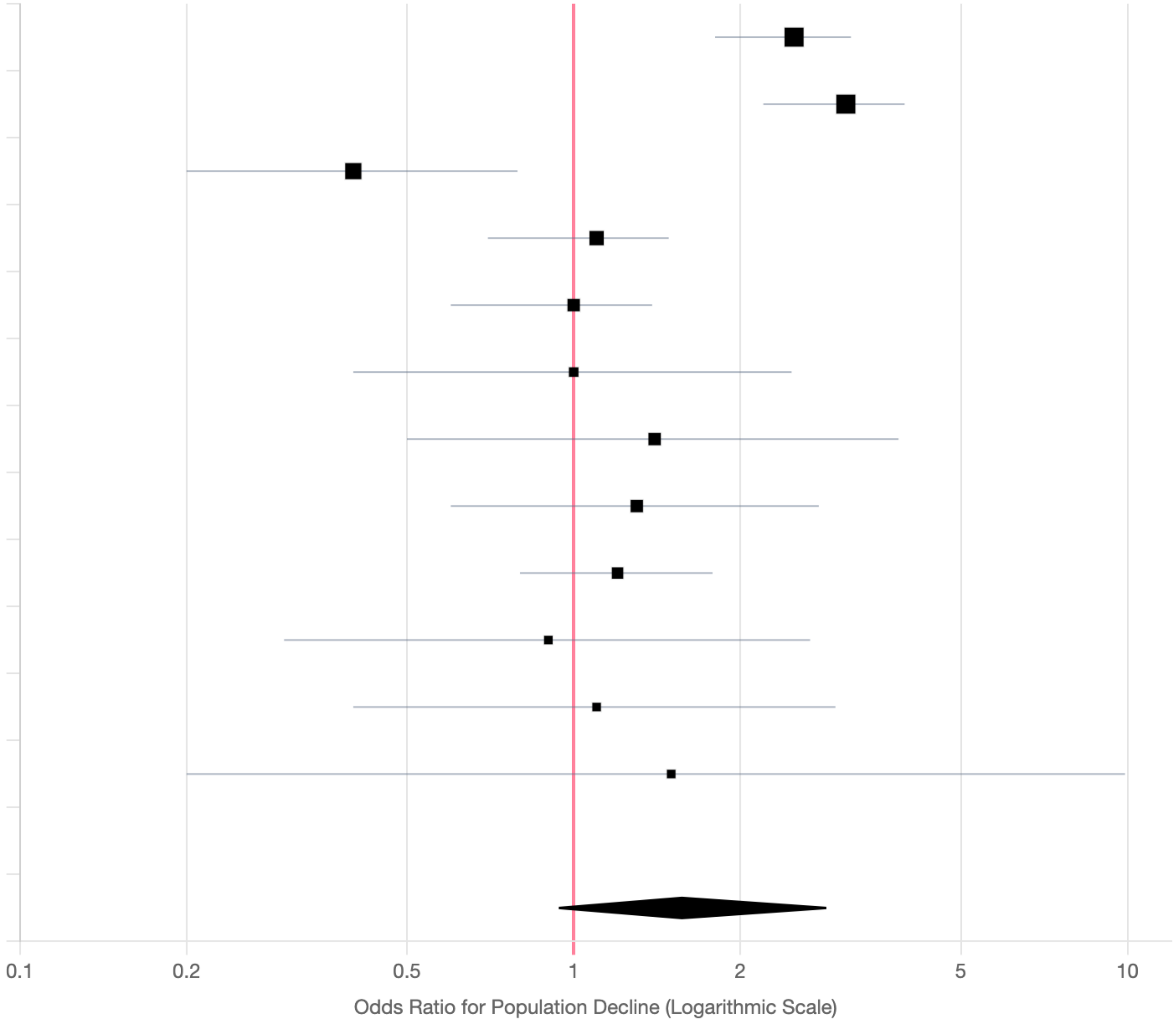


Effect size > 1 indicates an increased risk of population decline
